# Cross-Species Evaluation of qPCR Reference Genes in *Diabrotica speciosa* (Coleoptera: Chrysomelidae) using Diabrotica virgifera virgifera sequences

**DOI:** 10.1101/2023.12.05.570222

**Authors:** Luiz Renato Rosa Leme de Souza, Nayara Costa de Carvalho S. Okumoto, Henrique Marques-Souza

**Affiliations:** Department of Biochemistry and Tissue Biology, Universidade Estadual de Campinas, Campinas, São Paulo, Brazil; Department of Bioprocess Engineering and Biotechnology, Universidade Federal do Paraná, Curitiba, Paraná, Brazil

## Abstract

The cucurbit beetle *Diabrotica speciosa* is one of the most important corn pests in Brazil, seen as responsible for about 10-13% losses in maze crops, especially in the southern region of Brazil. The development of sustainable technologies for specific pest control, such as RNAi, depends on the identification of target genes and on the characterization of the gene expression profile of these genes in different developmental stages. Gene expression analyses, on the other hand, depend on reliable reference genes for the normalization of RT-qPCR data. In this study, we performed a cross-species evaluation of qPCR reference genes already validated in *Diabrotica virgifera virgifera*, due to its relevance as an important corn pest in North America. The expression of the reference genes EF1α, β-actin, β-tubulin, and GAPDH was evaluated in different developmental stages of the *Diabrotica speciosa*: egg, neonate, 3^rd^ instar larva, pupa, and adult. The stability of the reference genes was analyzed using the ΔCt method, BestKeeper, GeNorm, and NormFinder algorithms available at RefFinder web-based tool. From all four genes, β-actin and EF1α were the most suitable for normalizing genes, with both expression and stability ranking position in accordance with the results reported for *Diabrotica virgifera virgifera*. Our findings provide valuable information for researchers studying gene expression during different developmental stages of this species and highlight the potential of validating cross-species reference genes.

## Introduction

Normalizing genes play a key role in various experiments, such as the use of RNA interference (RNAi) aimed at pest control, which involves silencing the gene of interest in the insect, causing it to die or become sterile. The cucurbit beetle *Diabrotica speciosa* (*D. speciosa*) (NCBI:txid133126) is a notorious agricultural pest in Brazil, causing significant economic losses in various crops, including soybeans, rice, maize, beans, and potatoes [1,2]. This beetle belongs to the Chrysomelidae family and is commonly known in Brazil as “Vaquinha”. Traditionally, the control of this pest is based on the use of chemical insecticides, which can have negative impacts on the environment and human health, leading to a growing need to develop alternative pest control strategies to mitigate the use of agrochemicals. The RNAi technology represents a specific and sustainable alternative for pest control, as it allows for the inhibition of proteins essential to the insect via specific RNA-induced gene silencing. To identify such targets, one must characterize the expression of essential genes in different stages and growing conditions the insect pest will experience in the field, using Real Time quantitative PCR (RT-qPCR).

RT-qPCR is a widely used technique for quantifying gene expression levels in various biological samples. It is based on the polymerase chain reaction (PCR) method, in which a DNA template is amplified using specific primers and a fluorescent dye to quantify the amount of amplified DNA. Its popularity in studying gene expression stems from its high sensitivity, precision, and reproducibility, with a critical aspect being the selection of the appropriate reference gene for data normalization. To be considered a reference, a gene must have constant expression levels in all samples analyzed, to adjust for the differences in the quality, quantity, and efficiency of RNA reverse transcription. Identifying these appropriate genes is fundamental to obtaining accurate and reliable gene expression data.

Several studies have investigated the suitability of different genes in various tissues and organisms, highlighting the need to consider the selection of reference genes for normalization carefully and that several reference genes are often required for accurate normalization [3–6]. However, to our knowledge, there is still no study on reference genes in *D. speciosa*, with only 65 nucleotide sequences deposited in the NCBI database at the time of this publication, and given the substantial implications of *D. speciosa* in Brazilian agriculture, the need for the establishment of robust tools and analytical methodologies for investigating molecular and biological processes in this insect becomes paramount. To address this need, one potential strategy is to leverage data from a related study focused on the validation of reference genes in the species *D. virgifera virgifera* ([NCBI:txid50390](https://www.ncbi.nlm.nih.gov/Taxonomy/Browser/wwwtax.cgi?id=50390&mode=info)) [7], whose transcriptome has been assembled [8] and to whom several molecular studies have been addressed [9–12] due to its importance as responsible for expressive losses in maze crops in North America.

In recent years, new tools have been developed to improve the accuracy and robustness of reference gene selection, such as RefFinder [13,14], which integrates commonly used algorithms and provides comprehensive rankings of the most stable reference genes. Many studies have applied these algorithms to select suitable reference genes in different biological systems, including plants, animals, and microorganisms [15–18], to guarantee the reliability and reproducibility of gene expression analysis [19]. Among the established algorithms widely adopted, are GeNorm [20–22], NormFinder [23–25], BestKeeper [26], and the ΔCt comparative method [27–29]. The GeNorm algorithm calculates the measure of gene expression stability (M) by analyzing the pairwise variation between all reference genes and selecting the genes with the lowest M values as the most stable reference genes. NormFinder is another commonly used algorithm that estimates the stability value of reference genes by evaluating intra- and inter-group variations in expression levels. BestKeeper calculates the coefficient of variation (CV) and standard deviation (SD) of the reference gene’s expression levels and selects the genes with the lowest CV and SD values as the most stable reference genes. Finally, comparative ΔCt methods calculate the relative expression levels of reference genes and compare their stability using the standard deviation of the ΔCt values.

Reference genes often used in gene expression analyses in insect species are EF1α, β-actin, β-tubulin, and GAPDH. These genes are considered “maintenance (housekeeping) genes” because they are involved in basic cellular processes that are necessary for the survival and function of the cell. EF1α (elongation factor 1 alpha) is involved in protein synthesis [30,31], β-actin (Beta-actin) is a component of the cytoskeleton [32], β-tubulin (Beta-tubulin) is involved in the assembly of microtubules [33] and GAPDH (Glyceraldehyde 3-phosphate dehydrogenase) is involved in glycolysis [34]. These genes have been widely used due to their relatively constant expression profile in different tissues, developmental stages, and environmental conditions. In this study, we characterized the expression dynamics of genes EF1α, β-actin, β-tubulin, and GAPDH in different developmental stages to define the first to our knowledge set of reference genes for *D. speciosa*.

## Materials and Methods

### Biological Samples and Stages

Adult *D. speciosa* beetles were purchased from Pragas.com and used to start insect breeding in our laboratory to obtain different developmental stages (DDS). Three biological replicates of n individuals were collected for each stage analyzed: egg (n=150), neonate (n=50), 3^rd^ instar larva (n=20), pupa (n=5), and adult (n=4). Biological replicates were randomly collected from the breeding cage, immersed in liquid nitrogen, and forwarded to RNA extraction.

### Primer sequences

The primer sequences were sourced from the reference study targeting the species *D. virgifera virgifera* [7], and sent to ThermoFisher scientific™ for primer synthesis.

### *D. speciosa* DDS RNA extraction and cDNA synthesis

RNA extraction was performed using TRIzol™ reagent, then analyzed quantitatively by Nanodrop®2000 spectrophotometer and qualitatively by Agarose Gel Electrophoresis (data not shown). Using the RNA extracted, cDNA was synthesized using the Applied Biosystems™ High-Capacity cDNA Reverse Transcription Kit on an Eppendorf® Mastercycler® Pro. After that, cDNA stock solutions were diluted to working solutions at a concentration of 100 ng×μL^-1^ and saved at -80 °C for RT-qPCR assays.

### RT-qPCR

RT-qPCR assays were performed to analyze primer efficiency (I) and candidate reference gene stability (II). For (I), a microplate FrameStar® 384 Well Skirted PCR plate was assembled with *D. speciosa* DDS cDNA-containing pool (serially diluted, 10-fold), *D. virgifera virgifera* specific primers for each candidate reference gene, and the qPCRBIO SyGreen Mix. The plate was carefully sealed, and the reaction was performed in a CFX384 Touch Real-Time PCR Detection System-BioRad at 95ºC for 2 min, 40x (95ºC for 5 s, 60ºC for 30 s) followed by a melt curve from 65 to 95ºC by an increase of 0.5ºC per 5 s. For (II), the same steps were followed, except for the cDNA template distribution. Instead of a pool, cDNA from DDS at the same concentration were carefully distributed on separate wells.

### Data Preprocessing and Statistics

Raw data were inspected for missing values (NaN) and outliers using Grubb’s test at 5% of significance (α=0.05) using the software Paste© 4.14 [35]. Once the RT-qPCR assay was performed using technical replicates (n=3) for each biological replicate (n=3), NaN and outliers (Grubb’s p < 0.05) were replaced by the average of the neighbor technical replicates. Ct distribution and the RefFinder analysis were performed using the average of technical replicates as input.

## Results

### Primers Specificity and Efficiency

The slope and R^2^ values of the standard curve were used to assess the efficiency and linearity of the qPCR assay. Our results showed an R^2^ of 0.9992, a slope of -3.419, and an efficiency (E) of 96.10% for EF1α primer sequence. For β-Actin primer sequence, the R^2^ was 0.9945, the slope was -3.9945, and E was 95.83%. For GAPDH primer sequence, the R2 was 0.9992, the slope was -3.365, and the E was 98.22%. For β-Tubulin primer sequence, the R^2^ was 0.9993, the slope was -3.573, and E was 90.48% (Fig 1). Furthermore, each qPCR primer pair produced a single expressive melting peak, (Fig 2).

**Fig 1.**
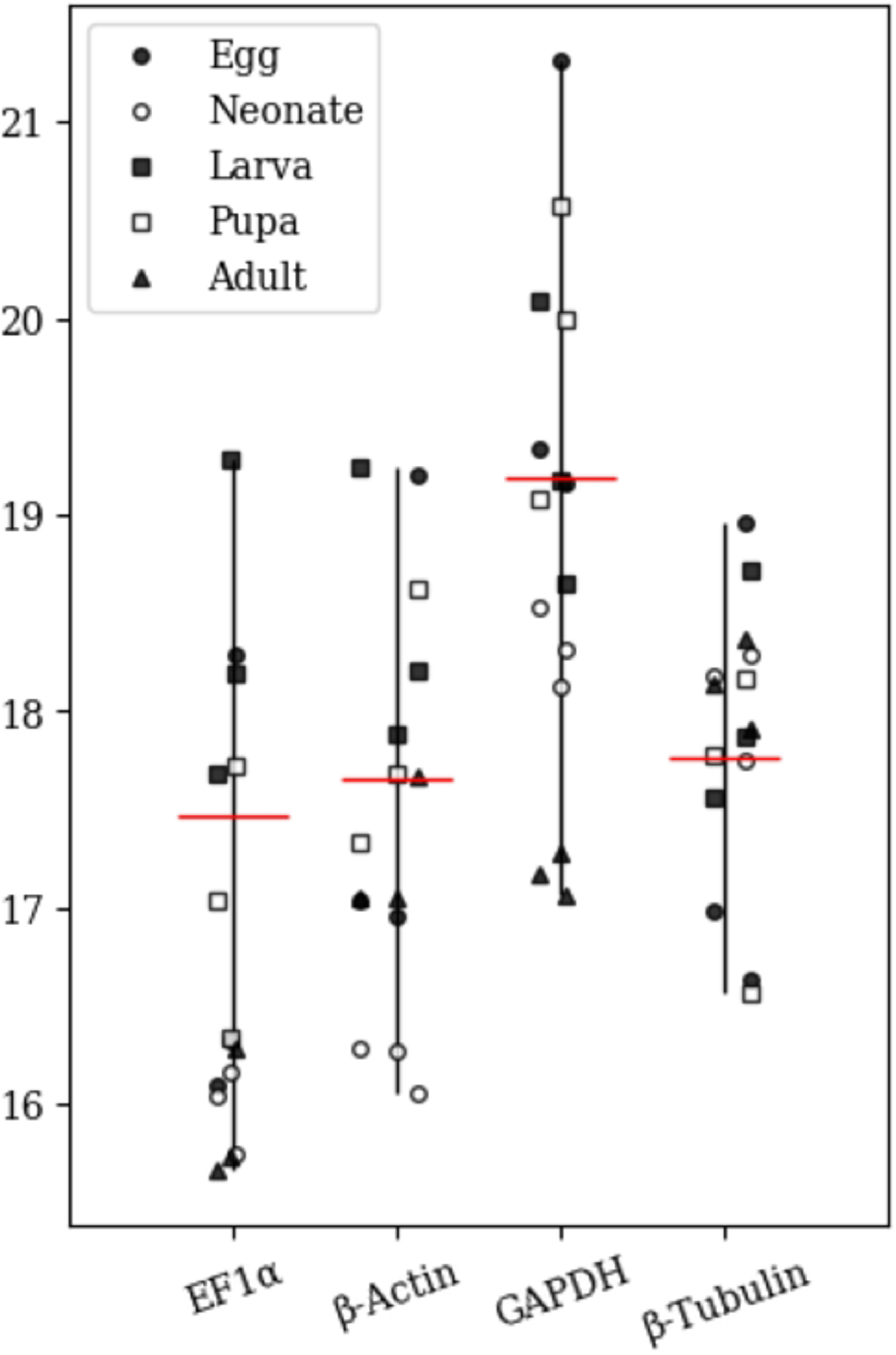
RT-qPCR standard curves for reference genes.

**Fig 2.**
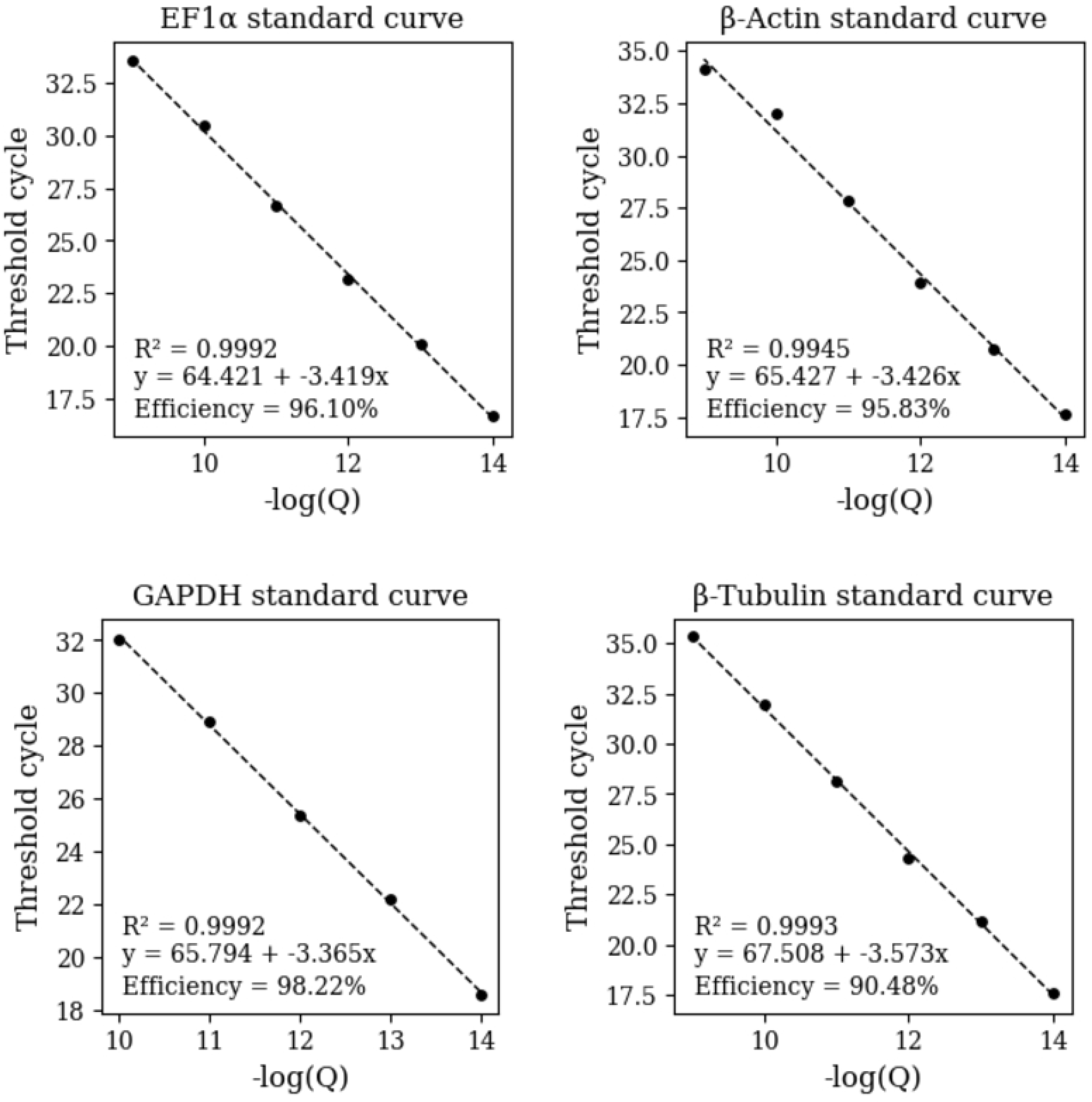
RT-qPCR melting peaks for reference genes.

### Cycle Threshold (Ct) Values

The data distribution and midrange (average between the highest and lowest values) of Ct were assessed for each gene (Fig 3). The gene EF1α midrange was 17.47, β-Actin was 17.65, GAPDH was 19.18, and β-Tubulin was 17.76, thus the EF1α was shown to be most expressed compared to the others, followed by β-Actin, β-Tubulin and GAPDH since the cycle threshold is inversely proportional to the expression level of the gene.

**Fig 3.**
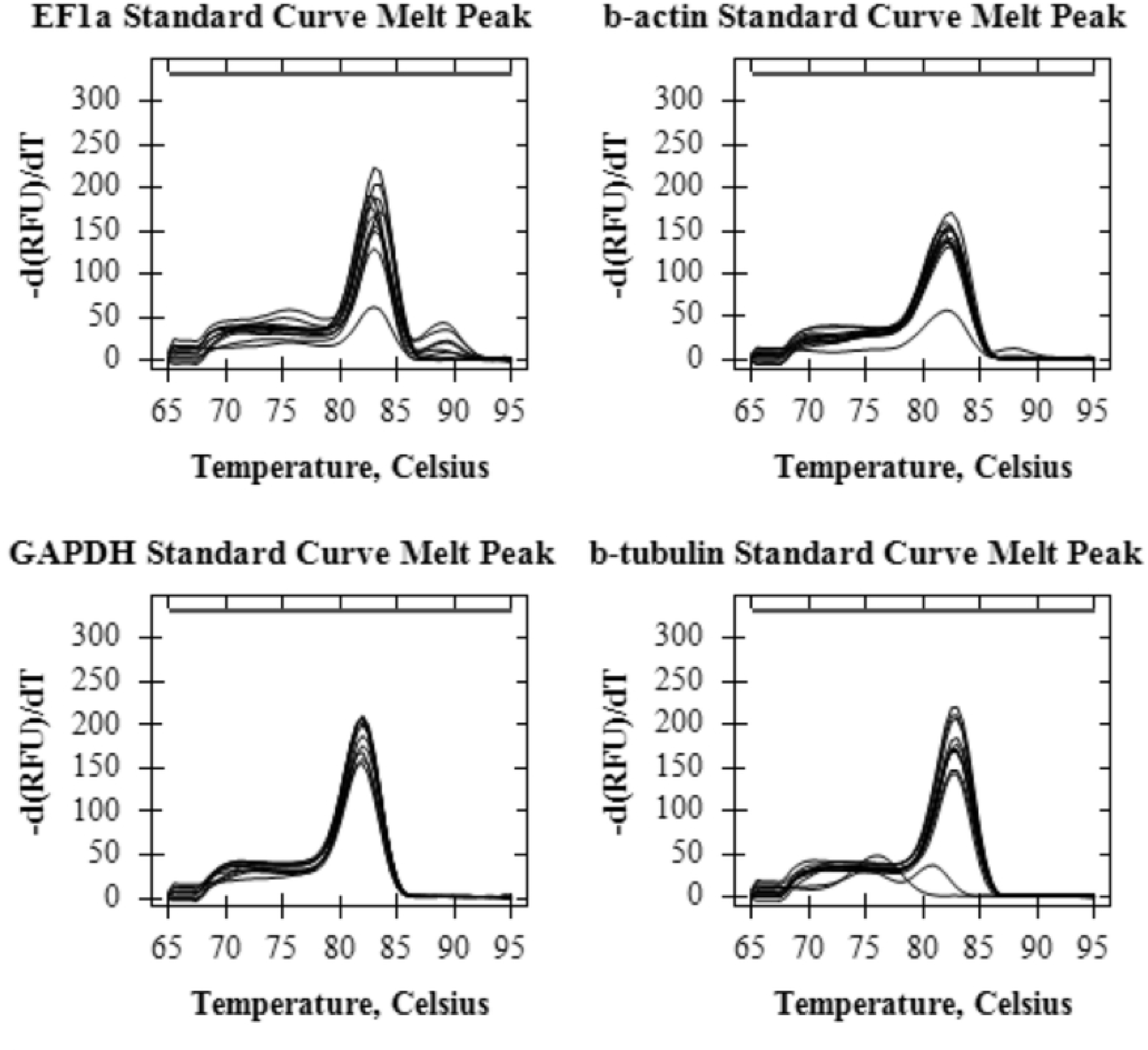
Cycle threshold (Ct) data distribution.

### Stability Analyses

GeNorm is a widely used algorithm that calculates the expression stability (average M value) of reference genes based on the average pairwise variation between them. According to this algorithm, the average M value for each gene was: 0.499 for β-Actin, 0.499 for EF1α, 0.773 for GAPDH, and 0.945 for β-Tubulin, thus resulting in the following ranking of stability: β-Actin = EF1α > GAPDH > β-Tubulin since the M value is inversely proportional to stability (Table 1).

**Table 1.** Stability values and ranking positions predicted by each algorithm.

| Gene name | Stability value "M" (GeNorm) | Ranking order (GeNorm) | Stability value (NormFinder) | Ranking order (NormFinder) | Stability value "SD" (Bestkeeper) | Ranking order (Bestkeeper) | Stability value (Delta-Ct) | Ranking order (Delta-Ct) | Stability value (comprehensive) | Ranking order |
| --- | --- | --- | --- | --- | --- | --- | --- | --- | --- | --- |
| β-Actin | 0.499 | 1 | 0.253 | 1 | 0.799 | 2 | 0.797 | 1 | 1.189 | 1 |
| EF1α | 0.499 | 1 | 0.323 | 2 | 0.955 | 3 | 0.809 | 2 | 1.861 | 2 |
| GAPDH | 0.773 | 3 | 0.886 | 3 | 0.978 | 4 | 1.057 | 3 | 2.828 | 4 |
| **β-Tubulin** | 0.945 | 4 | 0.978 | 4 | 0.518 | 1 | 1.116 | 4 | 3.224 | 3 |

RefFinder output for each algorithm presenting the stability value considered by each one, and the respective ranking position of each candidate reference gene.

The ΔCt method is calculates the relative expression of reference genes and compares all the samples in pairs. According to this method, the stability value for each gene was: 0.797 for β-Actin, 0.809 for EF1α, 1.057 for GAPDH, and 1.116 for β-Tubulin, thus resulting in the following ranking of stability: β-Actin > EF1α > GAPDH > β-Tubulin, (Table 1). BestKeeper is a statistical algorithm that evaluates the expression stability of reference genes based on the standard deviation (SD) and coefficient of variation (CV) of their Ct values. According to this algorithm, the stability value for each gene was: 0.799 for β-Actin, 0.955 for EF1α, 0.978 for GAPDH, and 0.518 for β-Tubulin, thus resulting in the following ranking of stability: β-Tubulin > β-Actin > EF1α > GAPDH, (Table 1). NormFinder is a statistical algorithm that uses a model-based approach to evaluate the expression stability of reference genes based on their intra- and inter-group variations. According to this algorithm, the stability value for each gene was: 0.253 for β-Actin, 0.323 for EF1α, 0.886 for GAPDH, and 0.978 for β-Tubulin, thus resulting in the following ranking of stability: β-Actin > EF1α > GAPDH > β-Tubulin, (Table 1).

## Discussion

In this study, we performed a cross-species evaluation of qPCR reference genes already validated in *D. virgifera virgifera* to allow for a direct use of these sequences as reference genes also in the species *D. speciosa*. Primers for the *D. virgifera virgifera* reference genes EF1α, β-actin, β-tubulin, and GAPDH was evaluated in different developmental stages of the *D. speciosa*. All primer pairs tested had slopes between -3.3 and -3.6, indicating an efficiency of 90-100%. The R^2^ values ranged from 0.98 to 1.0, indicating high linearity of the qPCR assay. Furthermore, a single expressive melting peak was obtained for each primer pair, indicating the high specificity of the PCR reaction. These findings support the notion that homology between both beetle species *D. virgifera virgifera* and *D. speciosa* is a path to be explored by further studies aiming at understanding this relation and its potential advantages. While cloning the specific gene in the studied species and design species-specific primer will always be the safer approach to define reference genes in any given species, the approach presented here demonstrate that primers already established in related species can allow for gene expression analyzes in situations where molecular cloning and DNA sequencing are limiting. To validate our cross-species qPCR evaluation approach, several algorithms have been employed to ensure reliable and accurate normalization. Among these algorithms, GeNorm stands out as a reliable and accurate tool for identifying stable reference genes in diverse qPCR experiments, recommending the use of top-ranked reference genes, as this is crucial for precise gene expression analysis [22]. The ΔCt method simplifies the process by calculating the relative expression of reference genes and comparing all the samples in pairs. The results are then ranked based on the standard deviation of the Ct values obtained [28,29]. Furthermore, the BestKeeper algorithm offers an alternative strategy. It analyzes and ranks the most stable reference gene(s) by considering standard deviation (SD) and coefficient of variation (CV) values. BestKeeper has proven to be a robust algorithm capable of handling a wide range of experimental conditions, making it reliable for identifying stable reference genes in various qPCR experiments [26]. NormFinder is a highly accurate algorithm well-suited for a comprehensive assessment of reference gene stability by considering the intra- and inter-group variations. It consistently demonstrates its reliability in identifying stable reference genes in a variety of qPCR experiments [25]. In summary, researchers can choose from a range of established algorithms to identify and prioritize the most stable reference genes for their specific qPCR experiments. This selection is a critical step in ensuring accurate and trustworthy gene expression analysis.

## Conclusion

The use of several algorithms to evaluate qPCR reference genes in a cross-species approach allowed for the identification of reference genes with high amplifying efficiency and linearity of the qPCR assays. This approach indicated the primer sequences for the genes β-actin and EF1α as the most reliable reference genes to normalize the Ct values resulting from RT-qPCR among different developmental stages of *D. speciosa* beetle (Coleoptera: Chrysomelidae). In addition, the melting peaks and efficiency values showed by the standard curves support the appliance of these primer sequences on molecular studies targeting this Diabrotica species. Furthermore, it reinforces the importance of using multiple methods to decide which genes to use as references based on a comprehensive approach.

## Acknowledgments

AI model ([OpenAI’s GPT-3.5](https://openai.com/)) and Grammarly were used to support the writing process of this paper. The authors acknowledge

LRRLS’s Fapesp Fellowship n: <u>2021/13976-2</u> and the Fapesp Research Funding n: <u>2018/23606-5</u>.

